# A stochastic model of corneal epithelium maintenance and recovery following perturbation

**DOI:** 10.1101/456947

**Authors:** E. Moraki, R. Grima, K. J. Painter

## Abstract

Various biological studies suggest that the corneal epithelium is maintained by active stem cells located in the limbus, the so-called Limbal Epithelial Stem Cell (LESC) hypothesis. While numerous mathematical models have been developed to describe corneal epithelium wound healing, only a few have explored the process of corneal epithelium homeostasis. In this paper we present a purposefully simple stochastic mathematical model based on a chemical master equation approach, with the aim of clarifying the main factors involved in the maintenance process. Model analysis provides a set of constraints on the numbers of stem cells, division rates, and the number of division cycles required to maintain a healthy corneal epithelium. In addition, our stochastic analysis reveals noise reduction as the epithelium approaches its homeostatic state, indicating robustness to noise. Finally, recovery is analysed in the context of perturbation scenarios.

## 1 Introduction

The cornea is the clear outer, avascular tissue that protects the eye’s anterior parts from inflammations and injuries (Watsky et al., 1995). Further, by controlling the light that passes through the eye it is estimated to contribute approximately 2/3 of the eye’s total focusing and optical power (Artal and Tabernero, 2008). The cornea, being a highly and organised set of different cell populations (Meek and Knupp, 2015), is arranged into five basic layers (Figure 1-(ii)): the corneal endothelium, the innermost layer keeping the corneal tissue clear (Oshima et al., 1998); a thin acellular layer known as Descemet’s Membrane; the stroma which covers nearly 90% of the cornea thickness (Kefalov, 2010); Bowman’s Layer, a transparent sheet of acellular tissue; finally, the outermost layer of the cornea, the epithelium, which accounts for approximately 10% of human cornea’s thickness (Reinstein et al., 2008) and varies according to species. Moreover, as the outward surface, the epithelium protects the eye from toxic UV irradiation (Marshall, 1985) and chemical injuries or pathological insults (Ruberti et al., 2011). It has been also characterised as “tight” (Liaw et al., 1992) since it has tight junctions and accounts for over 1/2 of the cornea’s total resistance to infection and fluid loss (Klyce, 1972).

**Fig. 1:**
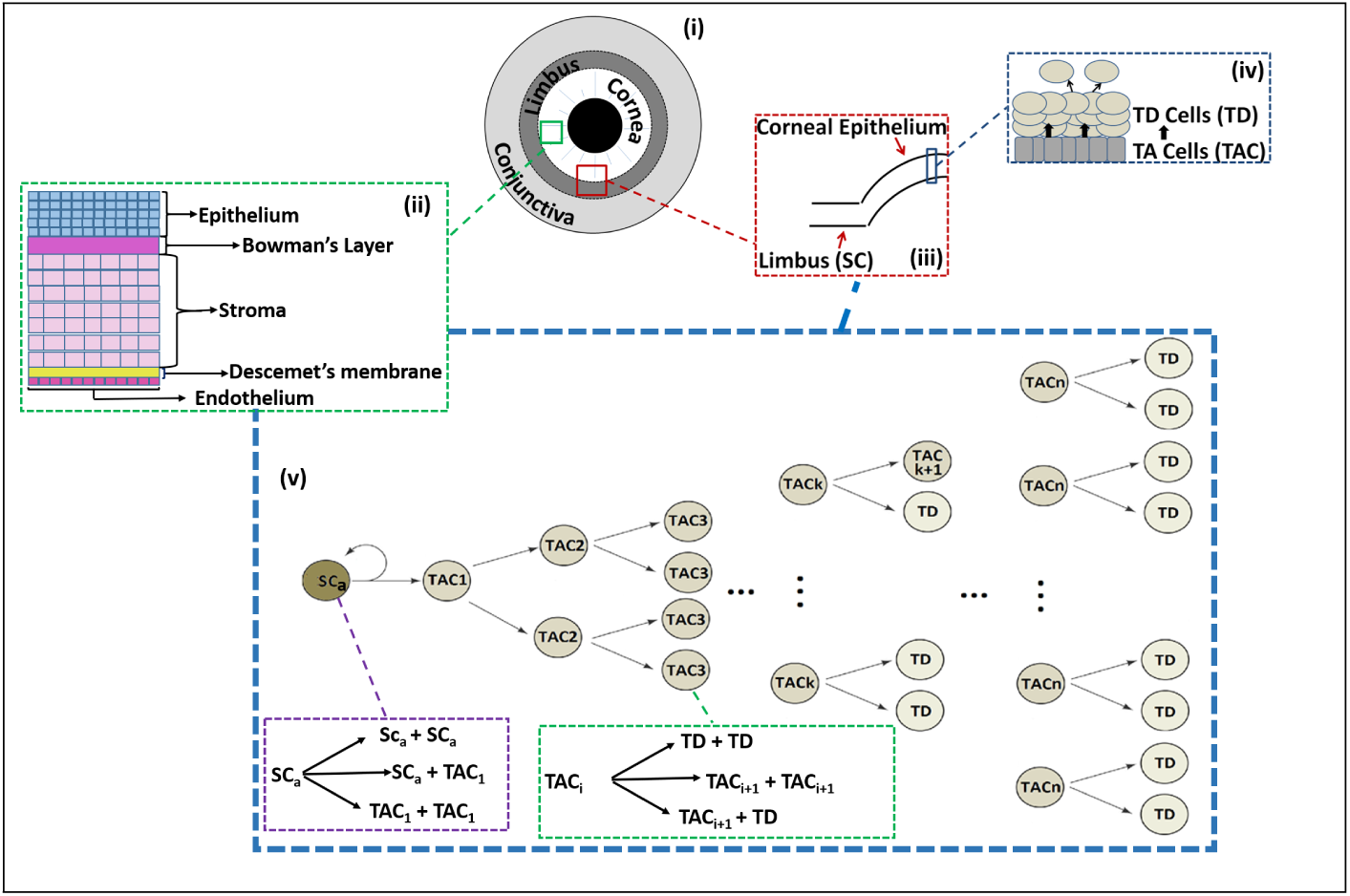
Biology of cornea stem cell maintenance. (i)-(iv) Surface and cross-section of the cornea, showing the limbal and central regions and the supposed positions of stem cells (SC), transient (or transit) amplifying cells (TAC) and terminally differentiated cells (TD). (v) Hypothesised model for stem cell maintenance, via active stem cell (*SC_a_*) division into multiple generations of TAC cells before eventual terminal differentiation (the production of TDs from cells of the last TAC generation, *n*).

The corneal epithelium is composed of 5−7 cell layers (Toropainen, 2007). The conventional view is that, during normal homeostasis, the corneal epithelium is maintained by limbal epithelial stem cells (SCs) that are located in the basal epithelial layer of the “limbus”, a ring-shaped transition zone between the cornea and conjunctiva. The SCs replace themselves and produce transient amplifying cells (TACs), which divide and move centripetally across the corneal radius to populate the basal layer of the corneal epithelium. The TACs also produce more differentiated cells (TDs), which move apically through the corneal epithelial layers and are shed from the surface (Zieske, 1994), (Figures 1(i), 1(iii), 1(v)). Despite some alternative proposals (e.g. Majo et al. (2008)), this hypothesis is the most widely accepted and is supported by almost 40 years of clinical observations and basic science (Davanger and Evensen, 1971; Tseng et al., 1989; Dua et al., 1994; Sun et al., 2010; Ahmad, 2012).

Prior to the realisation of SC maintenance of the corneal epithelium, the X, Y, Z hypothesis had been proposed by Thoft and Friend (1983). According to this, cell loss (Z) is balanced by (1) replacement from centripetal movement of peripheral corneal cell (Y) and (2) basal epithelial cell proliferation (X). This hypothesis and that of corneal outer layer self-maintenance by basal proliferation (Hanna and O’Brien, 1960) pre-dated modern understanding of stem cells, which are now known to be key for maintaining corneal tissue integrity (Daniels et al., 2001). Hence, the X, Y, Z hypothesis is somewhat updated by the Limbal Epithelial Stem Cell hypothesis (Dorà et al., 2015) by changing the definition of Y to the production of basal TACs by SCs. Active limbal stem cell (*SC_a_*) division generates two cells (Figure 1(v)) where each has the potential to remain a *SC_a_* (and stay in the limbus) or become a transient amplifying cell (TAC) that moves into the basal layer of the corneal epithelium periphery (Morrison and Kimble, 2006; Ebrahimi et al., 2009). Note that, at an individual level, stem cells do not necessarily divide asymmetrically: stem cells can divide symmetrically into either two stem cells or two TACs. As a whole, though, asymmetric division prevails to give rise to “population asymmetry” (Klein and Simons, 2011). The first generation of TACs (*TAC*_1_) produced from stem cells proceed through their cell cycle, subsequently undergoing a symmetric or asymetric division (Figure 1(v)) into either two *TAC*_2_ cells, a *TAC*_2_ and a *TD* cell or two *TD* cells. The same procedure applies in subsequent TAC generations. Note, however, that evidence suggests TACs more frequently undergo symmetric divisions than asymmetric ones (Beebe and Masters, 1996). Once a TAC cell loses its self-renewal ability, it simply divides into two TD cells (Figure 1(v)). TD cells lose contact with the basal layer and move up through the epithelium until they are eventually shed from the surface (Figure 1(iv)). This proliferation process is believed to provide the necessary cells required to maintain epithelial homeostasis.

Clearly, the above suggests that there must be a sufficient number of *SC_a_*s in the limbus to maintain the corneal epithelium. If the number of *SC_a_*s decreases (e.g. surgery, injury, disease) the corneal epithelium could lose its capacity for homeostasis, eventually resulting in a corneal disease known as limbal epithelial stem cell deficiency (LSCD) (Chan et al., 2015). Observations on human and mouse corneal epithelia suggest that the number of coherent clones of active SCs that are capable of maintaining the corneal epithelium decreases with age. While more investigation is required, this reduction can be caused either by an increase in the proportion of quiescent SCs (*SC_q_*) or loss of active SCs (*SC_a_*) in the limbal area (Mort et al., 2012). Experimental studies have shown that LSCD is associated with conjunctivalization, vascularization and chronic inflammation of the corneal epithelium (Chen and Tseng, 1990; Kruse et al., 1990; Chen and Tseng, 1991) with conjunctival epithelial in-growth (conjuctivalization) the most reliable diagnostic sign of LSCD (Puangsricharern and Tseng, 1995). There are many known genetic and hereditary causes of LSCD (Puangsricharern and Tseng, 1995; Espana et al., 2002) which results in pain and chronic ocular surface discomfort. In particular, corneal conjunctivalization leads to loss of corneal clarity, making LSCD an extremely painful and potentially blinding disease (Ahmad, 2012).

In this paper, we use mathematical modelling to determine the constraints placed on the proliferation process for healthy maintenance of the epithelium. Specifically, we investigate the number of active SCs (*SC_a_*), division rates and maximum cycle number needed to maintain the basal corneal epithelium with sufficient TAC cells. To account for potential variability, we formulate a stochastic model for this proliferation process and subsequently derive the corresponding system of ordinary differential equations (ODE model) that describes the average behaviour. Inevitably, corneal epithelium integrity is crucial for vision and perturbations (such as wounds) can cause integrity loss, so we also investigate recovery after various perturbations.

This paper is structured as follows. In Section 2 we briefly review existing mathematical models before providing a detailed description of our model. In Section 3 a steady state and stability analysis is performed with the aim of obtaining the constraints on the proliferation process that ensures the integrity of the tissue is not compromised. Aiming to investigate the noisiness of the system, we calculate the second moments of the stochastic model via the Lyapunov equation, before a Fano Factor and Coefficient of Variation estimation is presented. In Section 4, perturbation scenarios are considered. Finally, Section 5 summarises the main results of this work and describes future extensions.

## 2 Mathematical Modelling

### 2.1 Brief Review of Existing Models

A sizeable literature has focused on corneal epithelium modelling, with the specific aim of describing wound healing (Sherratt and Murray, 1990, 1991, 1992; Dale et al., 1994a,b; Sheardown and Cheng, 1996). Several attempts have also been made to explore stem cell population dynamics within other tissues, for example, in the colonic crypt (Paulus et al., 1992, 1993; Meineke et al., 2001; Gerike et al., 1998). A number of models have focused on cancerous stem cell dynamics, including the computational model by Meineke et al. (2001) and the deterministic models by Boman et al. (2001) and Johnston et al. (2007). In Marciniak-Czochra et al. (2009) a three multi-compartment model was proposed to describe the proliferation and asymmetric division of SCs during hematopoiesis. They investigated three different possible regulation mechanisms through feedback signalling and indicated that external regulation of SC self-renewal rate is necessary. Alarcon et al. (2011) proposed a state-dependent delay differential equation model for the stem cells’ maturation process, proving global existence and uniqueness of solutions as well as existence of a unique positive steady state for which they compute its formula. They also propose examples of biological processes where their model could be applicable, specifically in the context of cancer. Rhee et al. (2015) proposed two computational approaches to explain the spiral patterns of TACs which can be seen in mosaic systems and proposed that spiral angles are stable in mature mouse corneas.

As far as we are aware, only three studies have specifically investigated corneal epithelium maintenance, focussing on the centripetal movement of epithelial cells. Sharma and Coles (1989) proposed a population balance model based on the X, Y, Z hypothesis to study the centripetal movement of epithelial cells and how these regenerated from stem cells located in the limbus, determining the centripetal migration rate of TACs. Second, a recent mathematical simulation model by Lobo et al. (2016) showed that when physiological cues from the rest of the cornea are absent, a centripetal growth pattern can develop from self-organised corneal epithelial cells. Using the same computational framework they extended this to study the origin and fate of stem cells during mouse embryogenesis and adult life. In addition, they proposed that population asymmetry and neutral drift (when a *SC_a_* is lost due to production of two TACs, it may be replaced by a neighbouring *SC_a_* producing two *SC_a_*s, with this replacement leading to stochastic neutral drift of *SC_a_* clones) result in SC clone loss over the lifespan. Moreover, they showed that cell movement towards regions of excess cell loss due to blinking is feasible (Richardson et al., 2017).

### 2.2 Formulation of the Model

Here we ask the basic question: what are the constraints on active SC (*SC_a_*) numbers, proliferation rates and generations required to maintain a healthy epithelium? To that end we construct a simple stochastic model based on an analogy to chemical reactions, allowing us to account for random fluctuations in the cell numbers. While the migration of cells within the corneal epithelium is undoubtedly important, a primary determinant of the number of TACs would be the proliferation kinetics and, consequently, our current investigation focuses solely on this aspect. Specifically, we consider the dynamics in the basal layer, effectively assuming that maintenance of this layer is the key to overall homeostasis of the epithelium. The details of our model are as follows:

1. The *SC_a_*s located in the limbal basal layer (considered a one-dimensional ring, but we ignore spatial considerations in the present formulation) are assumed to divide with rate *α* into either: (a) two TACs of the first generation (*TAC*_1_) with probability *q_T,T_*; (b) two *SC_a_*s with probability *q_S,S_*; or (c) a *SC_a_* and a *TAC*_1_ with probability *q_S,T_* = 1 − *q_S,S_* − *q_T,T_* (see Figure 2a). Note that for the *SC_a_* division rate (Figure 2a) we have 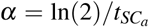, where 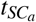 is the mean *SC_a_* doubling time, and
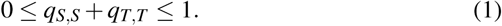 Across the lifespan of an organism, ageing is likely to result in declining *α* (Liu and Rando, 2011; Nalapareddy et al., 2017) and hence one could treat this parameter as a function of time. Moreover, *α*, *q_S,S_*, *q_T,T_* and *q_S,T_* are also likely to be functions of, amongst others, the available space to proliferate or chemical factors which allow feedback regulation of *SC_a_* division. In the interests of developing the simplest possible model we currently treat these parameters as constants. Note also that in the interests of model simplicity, we presently ignore either transitions of stem cells between quiescent and active states or potential reverse differentiations from TAC cells back into stem cells. Hence, the model is probably best viewed as operating over a relative short time span where such assumptions are reasonable. Note that a short time span refers to a timescale over which we can reasonably expect there not to be significant changes due, say, to aging of the organism. For example, in the case of a mouse, the order of a few months.
2. We denote by *TAC_i_* the *i^th^* TAC generation (where 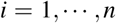). TACs located in the basal epithelial layer (considered to be a hemisphere of one-cell thickness) are assumed to divide with rate *β* into: (a) two TACs of the next generation (*TAC_i_*_+1_) with probability *p_T,T_*(*i*); (b) a *TAC_i_*_+1_ and a TD cell with probability *p_T,TD_*(*i*); or (c) two TD cells with probability *p_TD,TD_*(*i*) = 1− *p_T,T_* (*i*) − *p_T,TD_*(*i*) (see Figure 2b). Note that for the TAC division rate *β* (Figures 2b, 2c) we have *β* = ln(2)*/t_TAC_*, where *t_TAC_* is the mean TAC doubling time, and
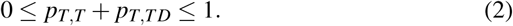 Similarly to *SC_a_*s division rates, ageing also causes TAC proliferation rates to decline (Liu and Rando, 2011; Nalapareddy et al., 2017) and hence *β* could also be considered a function of time. However, again we neglect this in the interests of simplicity. We do, however, assume that different TAC generations have different self-renewal abilities (Lehrer et al., 1998), assigning probabilities *p_T,T_* and *p_T,TD_* to be functions of the TAC generation *i*.
3. TAC cells of the very last generation (*TAC_n_*) are assumed to lose their self-renewal ability and division automatically leads to two TD cells as shown in Figure 2c. Note that one can theoretically set *n* = ∞ to give cells unlimited self-renewal capacity.
4. TD cells, once produced, lose contact with the basal layer of the epithelium, move up the layers and are eventually shed at a rate *γ* (Figure 2d).

**Fig. 2:**
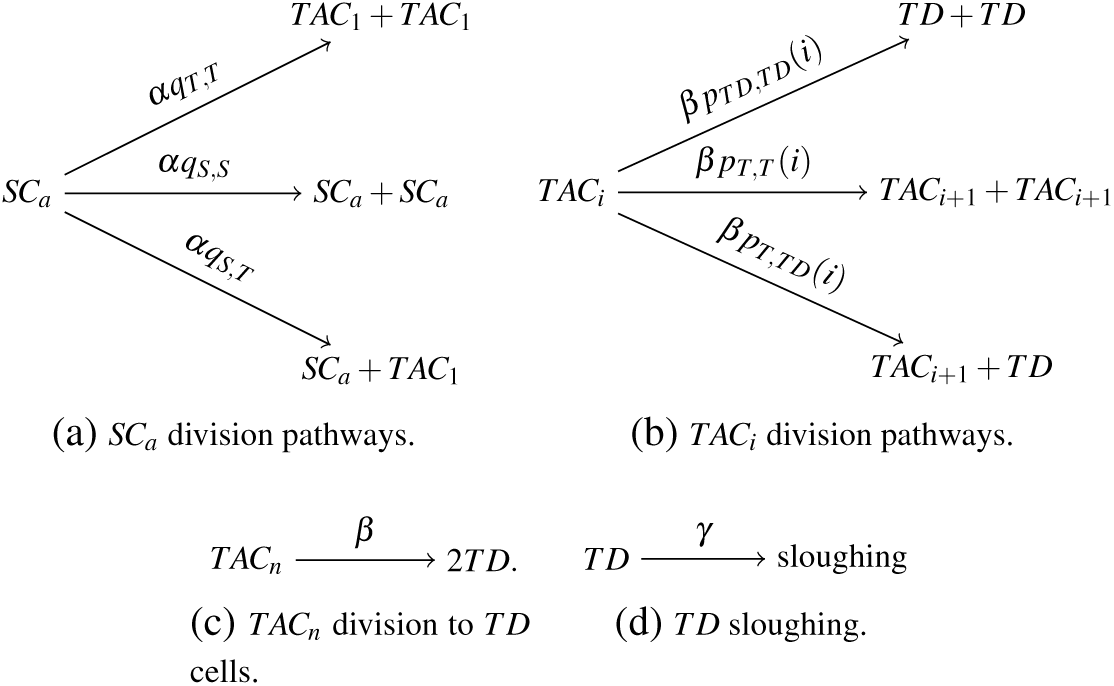
Possible *SC_a_* and TAC division pathways, and TD sloughing.

Given the above (1, 2, 3 and 4) one can write the chemical master equation (CME) using simple probabilistic laws (Gillespie, 1992). Let 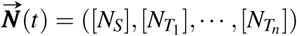 be the system’s composition vector, where [*N_S_*] is the *SC_a_* number and 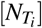 is the number of TAC cells in generation *i*, and *n* is the highest TAC generation number. Note that TD cells lose contact with the basal epithelial layer and, hence, are discarded from consideration. Let 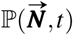 be the probability distribution for all possible states at time t, then the CME for our system reads

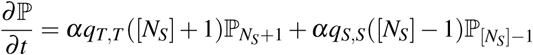

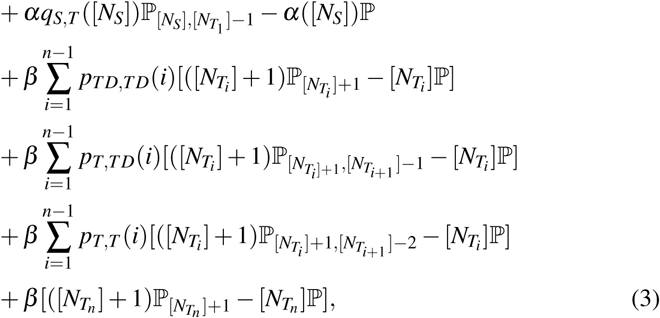

where 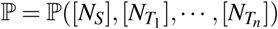, 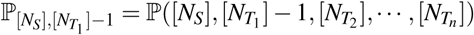, 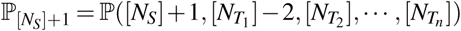, 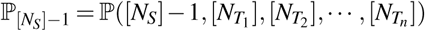,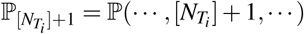, 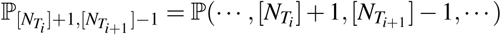 and 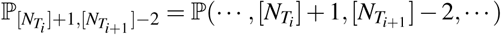

### 2.3 Derivation of Equations for the Mean Values

Having formulated the model via linear reactions, we can exploit the well known fact that the time evolutions for the stochastic mean values (the first moments of the Chemical Master Equation) are exactly equal to the solutions of the corresponding deterministic rate equations (e.g. see (Erban et al., 2007; Grima, 2010)). Hence, recall that denoting the cell numbers by [*SC_a_*] = *N_S_,* 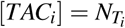 where *i* = 1, 2, …, *n* and [*TD*] = *N_TD_* and applying the Law of Mass Action, we obtain the following system of coupled ordinary differential equations (ODEs):
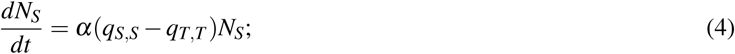

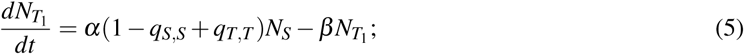

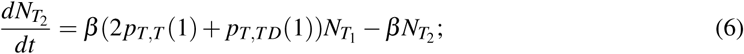

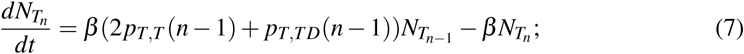

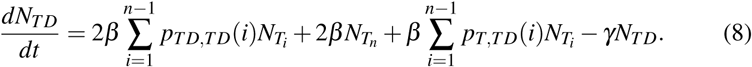

Note that *N_S_*, *N_TD_* and 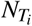 correspond to the *averages* for *SC_a_*, TD and TAC numbers, respectively 〈*N_S_*〉, 〈*N_TD_*〉 and 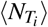. While the dynamics of TD cells do not impact on the dynamics of the stochastic model, and therefore have not been included in our original statement of the CME (Equation 3), it is a simple extension and we include their dynamics for completeness. Trivially we note that the mean *SC_a_* number remains constant if *q_S,S_* = *q_T,T_*, that is 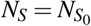 where 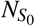 is the initial number of *SC_a_*s in the limbus. Further, Equation 8 decouples and can be ignored, allowing us to focus on system (4)-(7), provided it is assumed that TD cells cannot somehow return to a TAC state (see Figure 2d). Initial conditions will vary according to the context, for example with respect to whether we are exploring homeostasis or perturbation scenarios. We discuss these at the appropriate point and simply state that we close Equations 4-7 through some set of given initial conditions
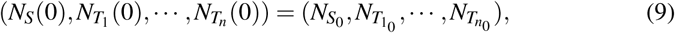

where, 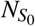, 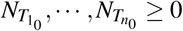.

Table 1 presents parameter ranges for the model; we refer to Appendix A for detailed discussion of values. Comprehensive understanding of the long time behaviour is vital to determine whether the corneal epithelium is maintained and we perform a steady state and stability analysis to address this. Specifically, we will determine the theoretical maximum number of TACs that the above model can generate (see Sub-section 3.1).

**Table 1:**
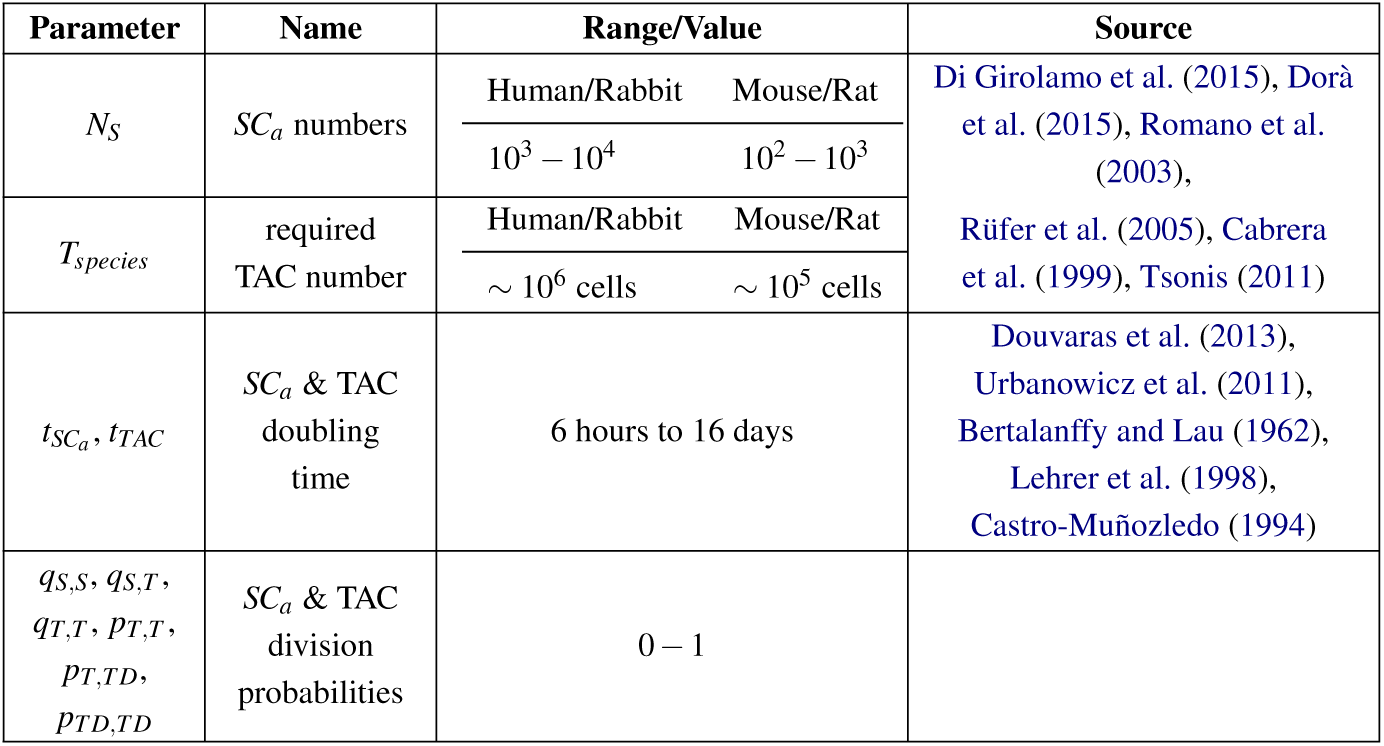
A list of all the principal parameters appearing in the models’ equations. The reader refers to the Appendix A for detailed reasoning behind the parameter choices.

## 3 Steady-State Analysis of the 1st Moment Equations

### 3.1 Maximum TAC Population

The tendency of stem cells to undergo asymmetric divisions is a common concept in the biological literature (Yoon et al., 2014). Symmetric divisions do not frequently take place and approximately half of the time they occur are self-renewing (Ebrahimi et al., 2009), suggesting *q_S,S_* ≈ *q_T,T_*. To simplify our models we therefore assume *q_S,S_* = *q_T,T_* and the average number of *SC_a_*s therefore remains constant.

To determine the densities (over the total corneal epithelial basal layer) of TAC generations that can be created by the model we use a straightforward steady state analysis. Specifically, we find a unique and stable steady state 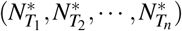 given by
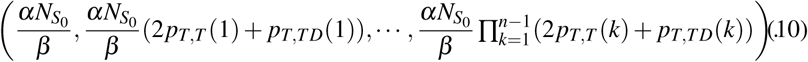

Moreover, explicit analytical solutions can be found to Equations 5-7. For example, the analytical solution for a total of two TAC generation is given by
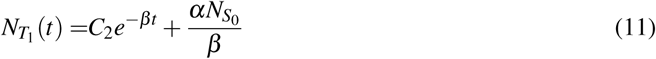

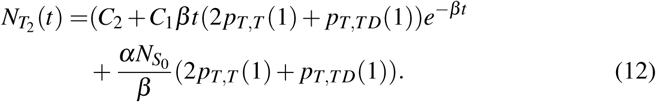

In general, for *n* TAC generations the solution for the *i*th TAC generation is given by
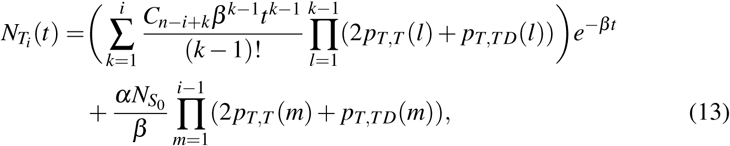

where *n* is the total number of the required TAC generations and *C_n_*_−*i*+*k*_ are the ODE integrating constants determined via the initial conditions.

As expected, the above shows that the number of TAC cells of generation *i* depends on the parameters of the proliferation process and the number of cells in the preceding generations. Note that the division rate of TACs (*β*) has a direct and clear effect on the rate of temporal dynamics. Taking the simple limit *t* → ∞ clearly shows solutions converge to the unique steady state solution (Equation 10).

Since solutions converge to Equation 10 we can interpret 10 as the number of each TAC generation that would be generated at homeostasis. Summing across all generations at steady state gives the total number of TAC cells (*T_SS_*) that can be generated at homeostasis:
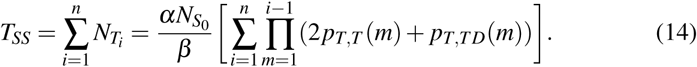

Note that 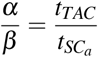 if division rates are expressed in terms of doubling times.

We suppose that the model is capable of generating and sustaining the corneal epithelium if the above exceeds the number of TAC cells required to fill a typical basal epithelium layer, which will of course vary with the size of the eye (and hence species). In other words, successful homeostatic capacity is subject to the condition
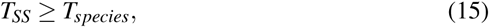

where *T_species_* refers to the total number of TAC cells that can be generated at homeostasis for different organisms. For example, based on typical basal cell and eye sizes for the mouse corneal epithelium we would require 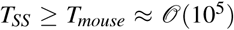 cells, while for human we could expect 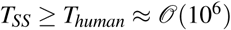 cells (see Table 1 and Appendix A).

### 3.2 Forms of TAC Division Probabilities

We consider two potential forms for the TAC division probabilities. Firstly, we take an analytically convenient step-function form
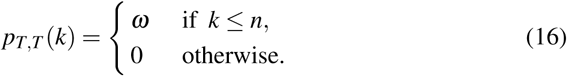

In the above 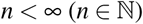 represents a strict upper limit on the number of divisions a TAC can make before automatically dividing into two TD cells. Obviously, *ω* being a probability implies 0 ≤ *ω* ≤ 1. The corresponding formulas for probabilities *p_T,TD_* and *p_TD,TD_* are
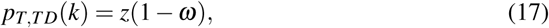

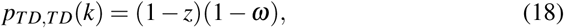

where constant *z* (0 ≤ *z* ≤ 1) represents the probability that a TAC undergoes an asymmetric division *if division into two TACs did not occur*. Note that the above ensures *p_T,T_* (*k*) + *p_T,TD_*(*k*) + *p_TD,TD_*(*k*) = 1.

Secondly, we consider the exponential form
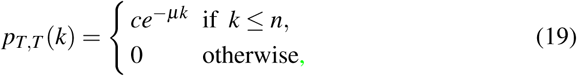

where *c* is constant which, for simplicity, we set *c* = 1. Note that choices *c* < 1 would result in earlier differentiation into TD cells and therefore act to reduce the total maximum number of TAC cells. We set the exponential decay rate *µ* > 0 and potentially allow *n* = ∞: theoretically, division from the *SC_a_* could lead to an infinite number of TAC generations but the probability exponentially decreases with generation number. This is in line with certain findings that TAC renewal ability decreases with the number of times they have divided (Yoon et al., 2014). The corresponding formulas for probabilities *p_T,TD_* and *p_TD,TD_* are of the form
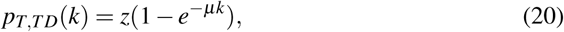

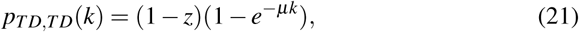

where, again, 0 ≤ *z* ≤ 1 represents the probability that a TAC undergoes an asymmetric division if division into two TACs does not occur. Again, it is easy to see *p_T,T_*(*k*)+ *p_T,TD_*(*k*)+ *p_TD,TD_*(*k*) = 1.

#### 3.2.1 Explicit Form for T_SS_ under Step-function Form

Substituting Equations 16 and 17 into Equation 14 we find
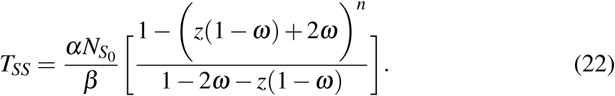

The “optimal scenario” demands *ω* = 1: TAC cells maximise their number by automatically undergoing symmetric divisions into two TACs of the next generation until terminal differentiation. In this case Equation 22 reduces to 
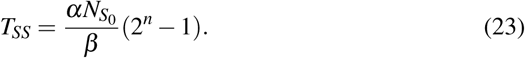

Thus, the number of TAC cells that can be generated increases with: (i) the number of *SC_a_*s; (ii) the number of TAC generations; and (iii) the ratio of *SC_a_*:*TAC* division rates. More so, this relationship allows us to generate parameter spaces for successful homeostasis, shown in Figure 3 based on benchmark figures for the size of a typical small epithelium, such as those of a mouse.

**Fig. 3:**
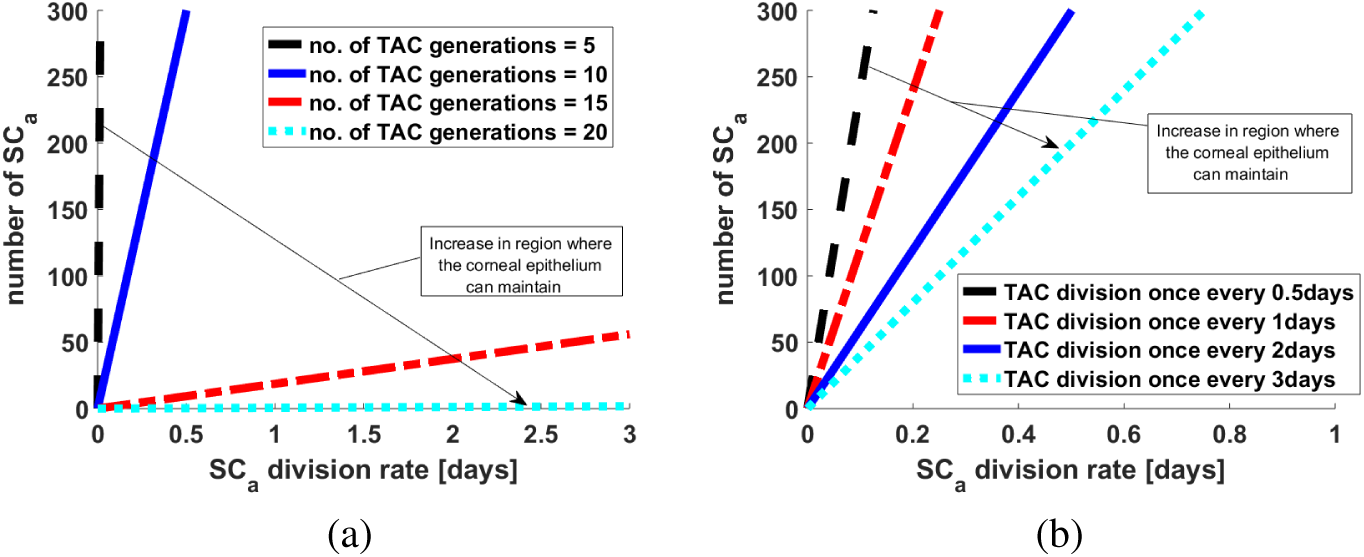
Parameter spaces in an “optimal case” (Equation 23), where only symmetric TAC divisions into two TACs are possible, showing significance of *SC_a_* number, *SC_a_* and TAC division rates and TAC generation number. The region of successful maintenance in the parameter space is the portion above each line, where “success” corresponds to generating the *>* 10^5^ basal epithelium cells required for a small (mouse-sized) cornea. (a) Change of the parameter space is shown for various maximum TAC generations, while fixing TAC division rates at once every 2 days. (b) Change of the parameter space is shown for a range of TAC division rates, while fixing the maximum number of TAC generations at 10.

***Parameter Spaces.*** We next expand to interspecies differences, exploring how parameter combinations would have to adapt to maintain a healthy corneal epithelium across eye sizes. Specifically, we consider two sizes: large (e.g. human/rabbit) and small (e.g. rat/mouse). In Figure 4a the successful parameter space region is shown in red for the large eye and in the union of red and blue regions for the small eye. Figure 4a(i) shows how the parameter spaces shift as the TAC doubling time increases, while Figure 4a(ii) illustrates how they change as *SC_a_* number increases. To provide more precise quantitative statements, consider the white dot in Figure 4a(i), which allows for a maximum of 8 TAC generations, a *SC_a_* doubling time of 2 days and 300 active stem cells. We see that a TAC doubling time of half a day would be insufficient to support either eye size, a doubling time of 2 days would be sufficient to support the small eye but not the large eye while a doubling time of 8 days would support both. Large TAC doubling times allows the TAC population to persist in the basal layer for longer, before eventual division into TD cells. In Figure 4a(ii) the dots represent a maximum of 8 TAC generations, and both TAC and *SC_a_* doubling times set at 2 days. Here we see that only 100 *SC_a_*s would be insufficient to support either eye size, 300 *SC_a_*s would support the small eye but not the large eye, while 1000 *SC_a_*s would support either eye size.

**Fig. 4:**
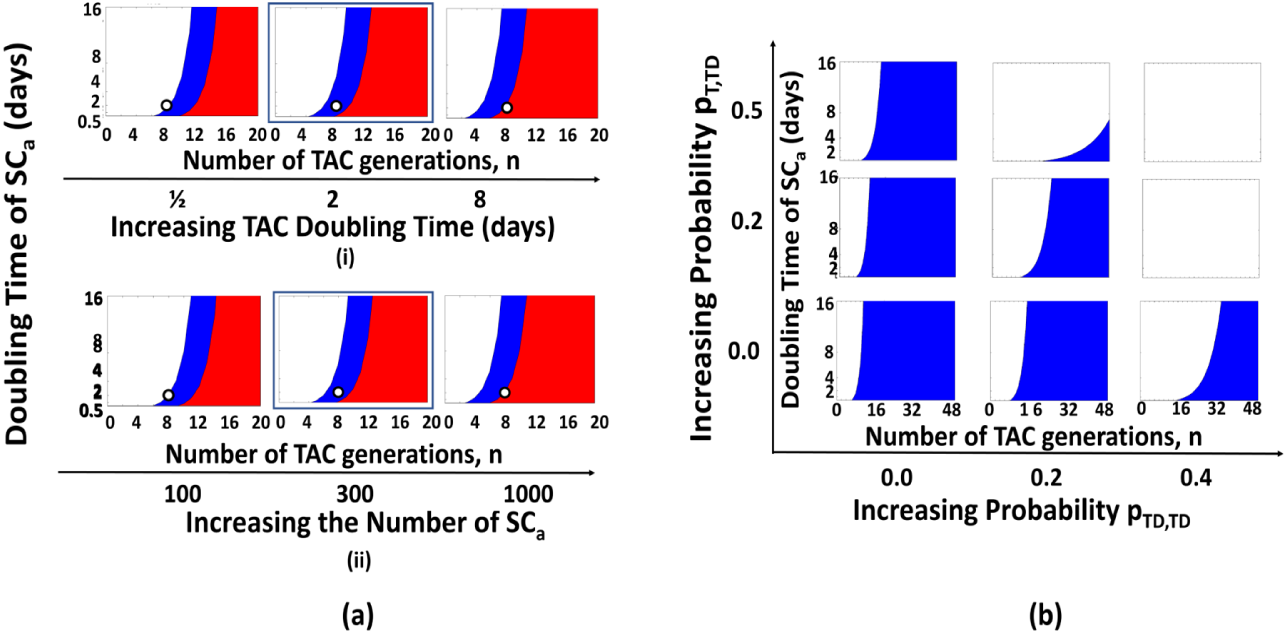
(a) Parameter space plots showing corneal epithelium maintenance demands for different eye sizes, under the “optimal scenario”. Red areas give parameter value ranges for maintenance in “large” corneas (i.e human/rabbit), while combined blue/red regions are for “small” corneas (i.e mouse/rat). Note that blue outlined figures specifically correspond to estimated parameter sets for a mouse cornea. *SC_a_* doubling time ranges between 1/2 to 16 days. (i) TAC doubling times of 1/2, 2 and 8 days and *SC_a_* number fixed at 300 cells. (ii) *SC_a_* number is 100, 300 and 1000 cells and TAC doubling time fixed at once every 2 days. (b) Parameter space plots under “sub-optimal” cases in which TAC assymetric division or premature terminal differentiation can occur. For these plots we fix the number of limbal *SC_a_* at 300, and TAC division rates at once every 2 days. Parameter spaces are plotted across the maximum permissible number of TAC generations and *SC_a_* division times for different *p_T,TD_*, *p_TD,TD_* combinations. White region shows where epithelium fails to maintain.

The plots in Figure 4a provide further visual insights into the parameter space for successful homeostasis: as expected from Equation 22, increases in TAC doubling time and *SC_a_* numbers lowers the required maximum number of TAC generations. Our main investigation will focus on mouse, since it is for this system that we have the most available data. Hence, considering the blue outlined frames, corresponding to a proposed normal scenario for mouse corneal epithelium where there exist roughly 300 *SC_a_*s (see Appendix A.1) and TAC cells divide once every two days (Urbanowicz et al., 2011), we see that somewhere between 5 − 12 TAC generations would be required for maintaining the mouse corneal epithelium (blue region) as we move across the range of *SC_a_* doubling times: fast stem cell divisions (once every 12 hours) would demand only 5 TAC generations, a longer doubling time (e.g. 16 days) would increase this to 12 TAC generations. Another TAC doubling time estimate is one every 3 days (Lehrer et al., 1998), and similar calculations would demand a TAC generation range of 4 − 11 according to the same range of *SC_a_* doubling times.

Moving beyond the optimal scenario, we next assume non-zero probabilities for asymmetric TAC divisions (division into a TAC and TD cell, *p_T,TD_*) and/or “premature terminal differentiation” (division into two TD cells before reaching the maximum generation, *p_TD,TD_*). Figure 4b shows the parameter spaces suggested by Equation (22) for a small eye scenario, as we progressively perturb probabilities *p_T,TD_* and *p_TD,TD_* from zero. Thus, the lower left most frame would correspond to parameter combination (*p_T,T_, p_T,TD_, p_TD,TD_*) = (1, 0, 0) while the upper right would correspond to (0.1, 0.5, 0.4). Increasing either *p_TD,TD_* or *p_T,TD_* from zero places a greater demand on the required number of TAC generations: these results follow naturally, since TAC cells prematurely enter the TD state.

Overall, while moderate *p_TD,TD_* and/or *p_T,TD_*, can be maintained, significant increases will result in a dramatic collapse in the size of the parameter space and unlikely maintenance. Note that, according to biological data TACs more often divide to cells of the same fate than asymmetrically (Beebe and Masters, 1996). Assuming a zero *p_T,TD_* probability the bottom line in Figure 4b suggests that the corneal epithelium can even be maintained when 6/10 times TACs divide to two TACs than two TDs, although there would be a significant increase in the number of generations required.

#### 3.2.2 Explicit Form for T_SS_ under Exponential Form

Next we consider our alternative generation-dependent division process, assuming the exponentially decreasing form. Specifically, we substitute Equations 19-20 into

Equation 14 to obtain
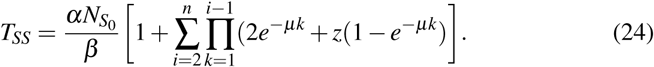

When no asymmetric divisions are possible (*z* = 0), Equation 24 becomes 
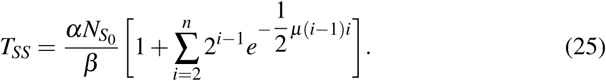

Note that by setting *µ* = 0 and *n <* ∞ we reduce to the step-function case analysed previously. When *n* = ∞ the parameter *µ* effectively replaces the concept of the maximum generations parameter: for small *µ* there is a high likelihood that TAC cells proceed through numerous generations before terminal differentiation, for large *µ* the reverse is true. Thus, small values of *µ* can generate high numbers of TAC cells at homeostasis and we focus on this parameter in subsequent investigations.

***Parameter Spaces.*** As Figure 5a shows, under symmetric TAC cell divisions (*z* = 0), decreases in either *SC_a_* number or the TAC doubling time demands smaller values of *µ* for healthy corneal maintenance. In Figure 5b(i) we show the relationship between *µ* and *n*, where we show that a small increase in *µ* can result in a substantially smaller parameter space.

**Fig. 5:**
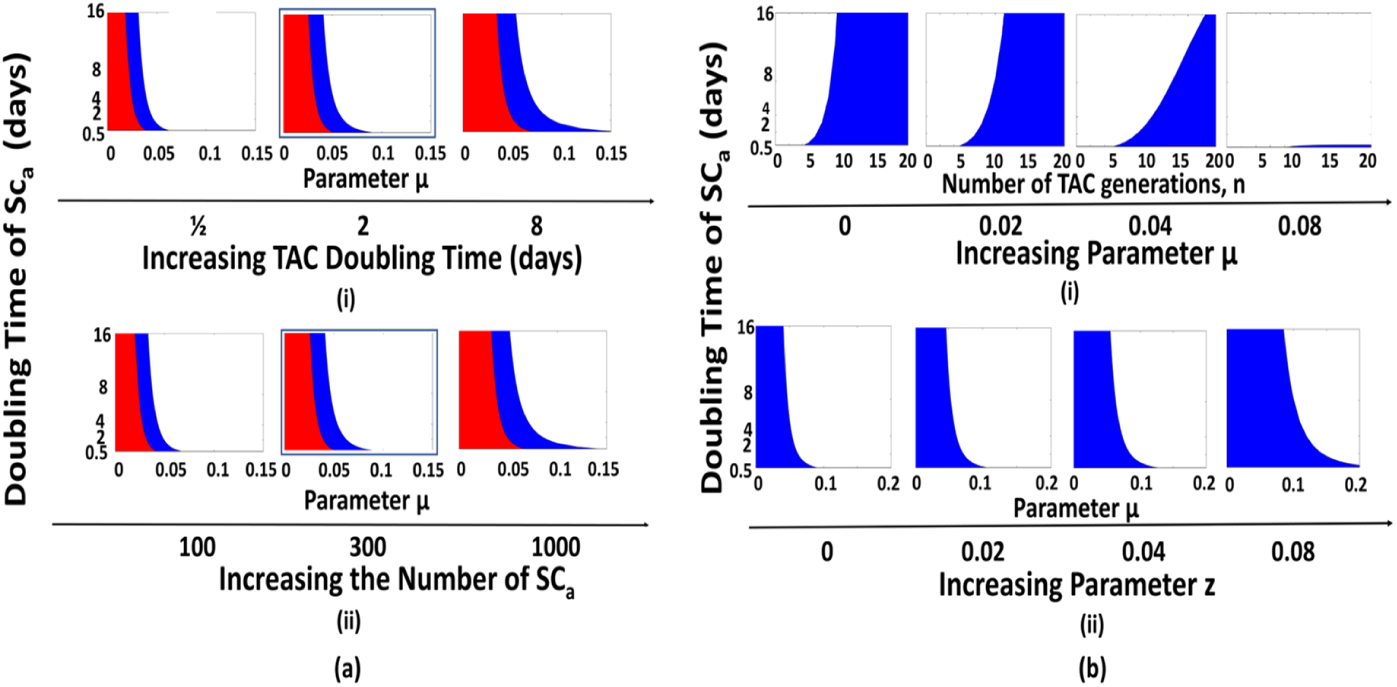
(a) Parameter space plots showing corneal epithelium maintenance demands for different eye sizes, for the exponential *T_SS_* form. Red areas give parameter value ranges for maintenance in “large” corneas (i.e human/rabbit), while combined blue/red regions are for “small” corneas (i.e mouse/rat). Note that blue outlined figures specifically correspond to estimated parameter sets for a mouse cornea. *SC_a_* doubling time ranges between 1/2 to 16 days. (i) TAC doubling times of 1/2, 2 and 8 days and *SC_a_* number fixed at 300 cells. (ii) *SC_a_* number is 100, 300 and 1000 cells and TAC doubling time fixed at once every 2 days. (b) Parameter space plots showing corneal epithelium maintenance demands for “small” eye size, for the exponential *T_SS_* form. For these plots we fix the number of limbal *SC_a_* at 300, and TAC division rates at once every 2 days. (i) Parameter space is plotted across the maximum permissible number of TAC generations and *SC_a_* division times for different parameter *µ* values. (ii) Parameter space is plotted across the maximum permissible value of parameter *µ* and *SC_a_* division times for different parameter *z* values.

Finally, in Figure 5b(ii) we extend to allow for asymmetric TAC division scenarios (*z >* 0) and investigate how the parameter space change. Under this scenario, a failure to divide into two next generation TAC cells does not automatically lead to two TD cells; rather, asymmetric divisions can allow a TAC cell to persist and hence the parameter space regime for successful maintenance is increased. Overall, however, we find that the results are generally consistent with those for the step function form implying model robustness with respect to the functional form, and therefore for the remainder of the paper we will use the step function form for its analytical convenience.

### 3.3 Derivation of Equations for the Second Moments

The steady-state analysis of the mean equations gives key insight into understanding the limitations of the average cell proliferation process, yet not the variability about the mean behaviour. Here we address this by deriving equations for the second moments at the unique steady state (Equation 10 in Subsection 3.1), using the CME (Equation 3).

Since the chemical system is monostable and composed only of first-order reactions, we can use the well-known fact that the second moments of the CME are given by the Lyapunov equation (Equation 26) (Schnoerr et al., 2017; Elf and Ehrenberg, 2003), where **C** is the correlation matrix 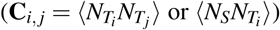,
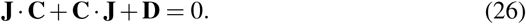

In the Lyapunov equation above (Equation 26), **J** is the Jacobian matrix and **D** is the diffusion matrix for the stoichiometries of the reactions along with their rates.

The diffusion matrix **D** is calculated by the stoichiometries of the reactions as described in stoichiometric matrix **S**, its conjugate transpose **S^T^** and the reaction rates which are described in the diagonal matrix **F** of the vector of macroscopic rates 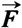:
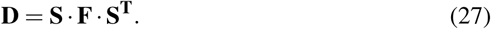

For calculation of matrices **J**, **S** and **F** included in Equations 26 and 27 from the chemical reactions we refer to Appendix B. Moreover, it is worth mentioning that the approach we use here is frequently used for Linear Noise Approximation (LNA) in biochemically reacting systems (Elf and Ehrenberg, 2003).

To obtain the correlation matrix **C**, we use a built-in Matlab function (lyap) for the Lyapunov Equation while changing the probabilities as to whether TACs are undergoing symmetric or asymmetric divisions. We use these results to investigate the stochastic properties of the system. In particular, we calculate the Fano Factor (FF) and the Coefficient of Variation (CV) in Subsection 3.3.1 (Thomas et al., 2013; Paulsson, 2005). The FF is a measure of how different are the second moments of the stochastic process, compared to those of a Poisson distribution with the same mean. The FF equals one for a simple birth-death process with constant rates. The CV is a measure of the size of the fluctuations relative to the mean; it is zero for a purely deterministic system.

#### 3.3.1 Noise

We use the Lyapunov Equation (Equation 26) to investigate how the variance in cell numbers differs from that of a Poisson distribution with the same mean by calculating the FF, defined as
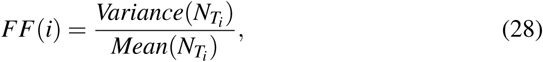

with *i* the TAC generation number.

To investigate whether FF increases (or decreases) as we move through TAC generations (and hence as more cells are added into the system) we numerically solve the Lyapunov equation (Equation 26) and, as a reference case, we fix the parameters for the number of *SC_a_* at 300, *SC_a_* and TAC division rate at once every two days and *p_T,T_* = 0.8 and *z* = 0.4. Moreover, we force a requirement to generate sufficient TAC cells to maintain a small corneal epithelium (10^5^ cells). The analysis shows that *FF* increases with the TAC generation. Specifically, our results show that while *TAC*_1_ cells follow a Poisson distribution (*FF* = 1), the distribution of cells in subsequent generations changes significantly to Super-Poissonian behaviour, since *FF >* 1 (Figure 6a). This shows that while for *TAC*_1_ cells the proliferation process is analogous to a simple “birth-death” process, this is not true for the subsequent TAC generations and variances in the cell densities will be larger than expected from Poisson statistics.

**Fig. 6:**
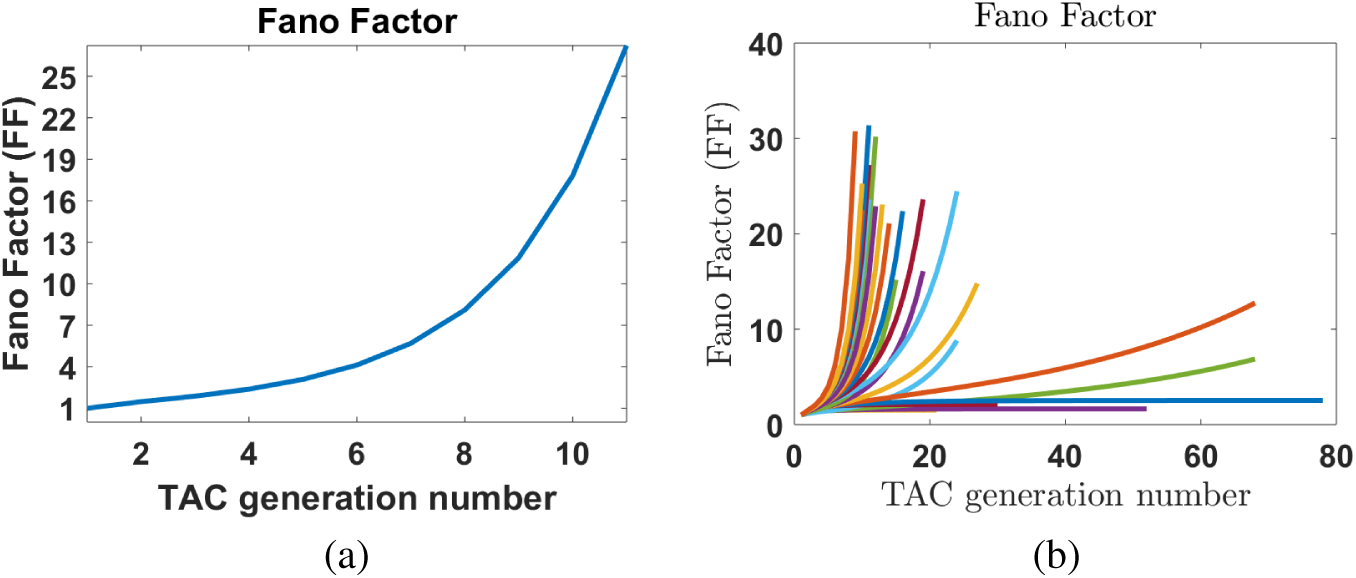
Plots show the increasing tendency of Fano Factor (FF) where *SC_a_* = 300, *α* = *β* = log2/2 and *T_tot_* = 10^5^ cells (similar to a small cornea epithelium of a mouse). (a) FF in reference case where *p_T,T_* = 0.8 and *z* = 0.4. (b) FF increases for all pairs of division probabilities (0.2 ≤ *p_T,T_* ≤ 1 and 0 ≤ *z* ≤ 1). Note that similar results concerning the FF increasing tendency are observed for all perturbation experiments about *SC_a_* numbers, *SC_a_* and *TAC* division rates.

To investigate if the FF increasing tendency was a result of parameter choice, we consider the ranges 0.1 ≤ *p_T,T_* ≤ 1 and 0 ≤ *z* ≤ 1. Simulations under these perturbations show similar results, with FF always increasing with the TAC generation number. Specifically, Figure 6b shows this FF increasing tendency which is either small or large. Small increases in FF as we move towards higher generations are reported when *p_T,T_* + *p_T,TD_ <* 0.5 (i.e. *p_T,T_* = 0.2 and 0 ≤ *z* ≤ 0.6) and hence, *p_TD,TD_ >* 0.5. This is logical as few TAC cells are generated in the epithelium and hence the variance in the cell numbers will be small. We recall that different pairs of *p_T,T_* and *z* result in different maximum TAC generation numbers, in order to generate the total of 10^5^ cells as shown in Figure 6b.

We then perturb first the *SC_a_* number in the limbus, followed by the *SC_a_* division rates and finally the TAC division rates, for a range of probabilities 0.1 ≤ *p_T,T_* ≤ 1 and 0 ≤ *z* ≤ 1. Note that for each experiment other parameters were fixed and set equal to those of our reference case. For all of these experiments, our simulations gave the same results (with respect to the increasing tendency of the FF) indicating that the FF increasing tendency is a persistent property. Our experiments above therefore indicate that the model is robust, in the sense of insensitivity to the chosen parameter values.

Next we study a second noise measure, the Coefficient of Variation (*CV*),
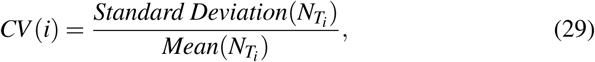

where *i* is the TAC generation number.

Larger *CV* implies a noisier system. In regions where the parameter combinations are capable of maintaining a small mouse corneal epithelium, the number of first generation TACs as they are pushed into the basal layer is slightly noisy, but noise rapidly decreases for higher generations. Since it is indeed the higher generations that contribute the bulk of the TAC cells, this suggests overall robustness of the system. As an illustrative example, consider 0.7 ≤ *p_T,T_* ≤ 1 and *z* = 0.4: the decreasing nature of CV with generation number is shown in Figure 7a. On the other hand, for a non maintaining region where *p_T,T_* < 0.4 (and *z* = 0.4) the CV increases with the number of TAC generations (see Figure 7b). In other words, non-robustness in the sense of noise is only observed in biologically unrealistic regimes, i.e. where the epithelium cannot be maintained.

**Fig. 7:**
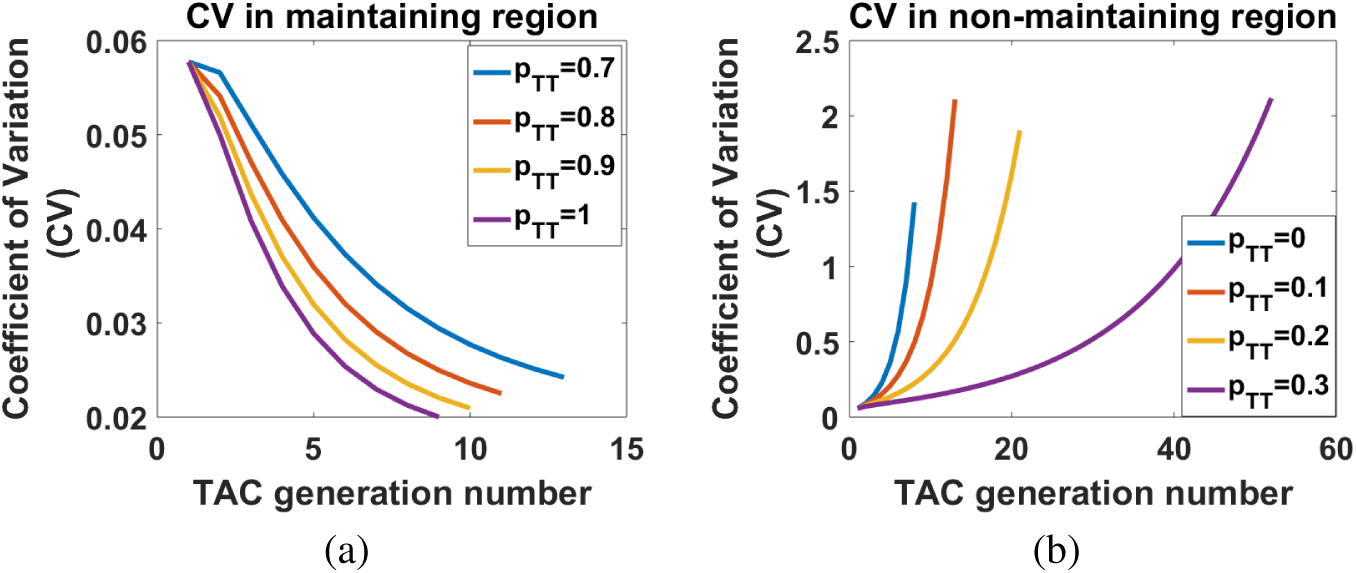
Plots show Coefficient of Variation (CV) in two different regions when *SC_a_* = 300, *SC_a_* and *TAC* division rates are set at once every two days and *z* = 0.4. (a) Maintaining region where 0.7 ≤ *p_T,T_* ≤ 1 and the CV decreases. (b) Non-maintaining region where 0 ≤ *p_T,T_* ≤ 0.3 and the CV increases.

To understand whether this is impacted by the TAC division probabilities, we solve as previously by fixing the total number of TAC that can fill the epithelium to be 10^5^, the *SC_a_* number in the limbus at 300 and the TAC division rate of once every 2 days. Over the full range of *SC_a_* division rates (Table 1), we find decreasing (increasing) noise for division probability pairs marked black (white) in Figure 8a. Note that the decreasing cases correspond exactly to the “biologically” relevant regime, i.e. those parameter combinations capable of sustaining a healthy cornea (Section 3.2).

**Fig. 8:**
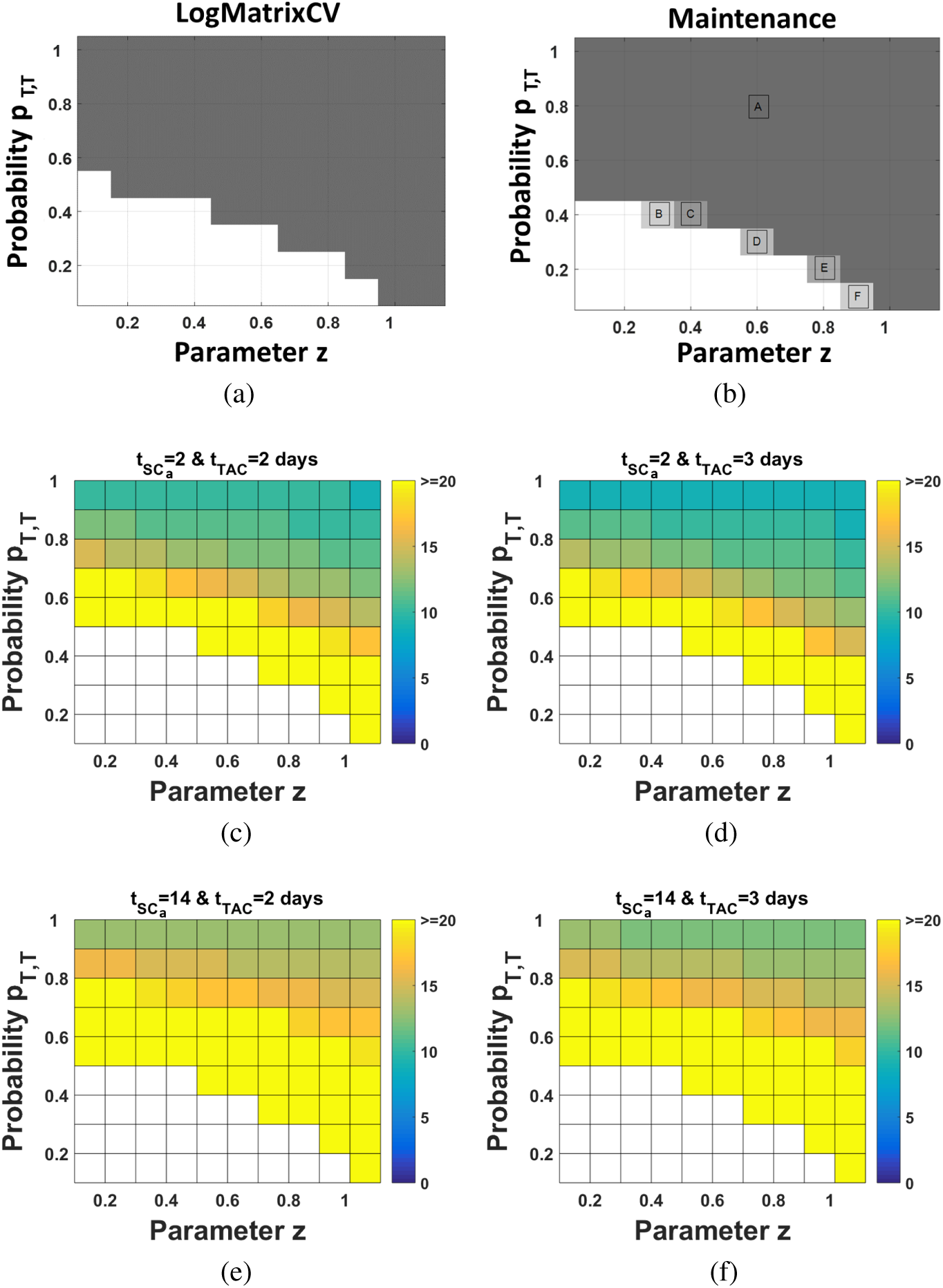
Noise decreases for the whole parameter space where corneal epithelium is maintained. (a) Coefficient of Variation. Black (white) area shows the parameter space where CV decreases (increases) with the increasing number of TAC generations. (b) Parameter space where corneal epithelium can maintain when 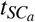 is from 6 hours to 14 days. Area A correspond to all pairs of *t_TAC_* and 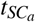. Area B to *t_TAC_* ≥ 8 *d* and 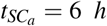. Area C to *t_TAC_* ≥ 2 *d* and 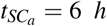, *t_TAC_* ≥ 4 *d* and 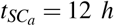 and *t_TAC_* ≥ 8 *d* and 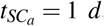. Area D to *t_TAC_* = 6,10 *d* and 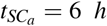 and to *t_TAC_* = 10 *d* and 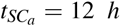. Area E to *t_TAC_* = 4, …, 10 *d* and 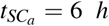 and to *t_TAC_* = 8, 10 *d* and 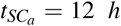. Area F to *t_TAC_* = 8, 10 *d* and 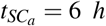. (c)-(f) TAC generations required for maintaining the epithelium when 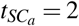, 14 days and *t_TAC_* = 2, 3 days. White area corresponds to parameter space not capable of maintaining the epithelium.

As a further test we altered the number of *SC_a_* in the limbus (100, 300 and 1000 cells) and again allowed *SC_a_* division rates to range from 6 hours to 10 days; results are shown in Figure 8a. Thus, *FF* always increases independently of the various parameter choices while *CV* decreases, but we can see a dependency on the division probability choices.

For further investigation into whether noise decreases for parameters at which the corneal epithelium is maintained we plot a *parameter space for maintenance* in Figure 8b. Here probabilities *p_T,T_* and *z* are perturbed (recall that *p_T,TD_* = *z*(1 − *p_T,T_*) and *p_TD,TD_* = (1−*z*)(1− *p_T,T_*); see Subsection 3.2). Moreover, *SC_a_* and TAC division rates are also perturbed, but such that the model generates a sufficient number of TAC cells to maintain a small corneal epithelium (10^5^ cells). Note, therefore, that since the TAC division rates vary we expect the TAC generation number to vary as well. Dark areas correspond to those parameter combinations which can generate 10^5^ cells, while those in white regions fail. Providing that parameters sit within *A* (biologically relevant, as already discussed in Subsection 3.2.1), a straightforward comparison between Figures 8a and 8b indicates that we have robustness to noise. It is only when the parameters sit on the threshold of the parameter space where we start to get potential noisiness of the system. Figure 8b suggests that for *p_T,T_* = 0.5 and 0 ≤ *z* < 0.1 the corneal epithelium is maintained (while noise increases in Figure 8a), but only for more than 20 TAC generations. For example, for *z* = 0 and 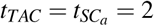 days the epithelium would require an unfeasibly large 334 generations. Note further that areas B-F correspond to TAC division rates of more than once every 3 days, beyond estimated values. Hence Figure 8a and 8b suggests that increasing noise only occurs under “abnormal” conditions. Figures 8c, 8d give the number of TAC generations required when the *SC_a_* doubling time is once every 2 days and the TAC doubling time is either once every 2 or 3 days (Lehrer et al., 1998). In Figures 8e and 8f the numbers of TAC generations required to maintain the epithelium are shown when 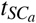 is of once every 14 days (an upper bound of the *SC_a_* division rate according to Douvaras et al. (2013)) and *t_TAC_* is set at once every 2 or 3 days respectively. When probability *p_T,T_* = 1 and 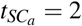 days, the required TAC generations are 9 and 10 for *t_TAC_* = 2 and 3 days respectively, while when 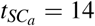 days the required generations are 12 and 11. This suggests that there is not a large difference in the required generations for large variations of 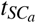 for biological estimated *t_TAC_* values.

Summarising, FF always increases with the numbers of TAC generations. While CV can increase, it only does so outside the relevant parameter region for maintenance. Hence, it is possible that parameters evolve over an individual’s growth into a state where noise reduces while maintaining the epithelium. Noise reduction would clearly be beneficial (Rao et al., 2002) for maintaining the epithelium, since fluctuations in cell numbers would be minimised. For a healthy individual we would expect the parameters to all sit comfortably within A, but a potential result in Limbal Stem Cell Deficiency would involve parameters (e.g. reduced *TAC* division rates) shifted towards the boundary, where one starts to “feel the effect” of noise.

## 4 Perturbation Experiments

### 4.1 Objectives

Here, we explore the behaviour under perturbations linked to pathological/wound healing type scenarios or biological experiments. For the in silico experiments we consider a specific reference parameter set, motivated by mouse cornea. Specifically, we set the *SC_a_* number to be 300 and their division rate once every two days (unless otherwise stated). With 9 TAC generations, a TAC division rate of once every 2 days, the step-function choice for TAC divisions and “optimal” division (*p_T,T_* = 1 and *z* = 0), this would enable a basal corneal epithelium to be supported of up to 153, 300 cells. For the remainder of the section our initial conditions are taken to be the homeostatic steady state distribution of TAC cells, and at some time *t_pert_* we apply some perturbation, where the exact form of perturbation will be defined at the appropriate point.

### 4.2 Pathological *SC_a_* Loss

First, consider the impact of pathological stem cell loss scenarios, defined as an *SC_a_* loss from the limbal area of the eye (e.g. as associated with LSCD or due to injury/experimental extraction). Note that here we do not consider any mechanisms that may boost *SC_a_*, e.g. via symmetrical divisions to two *SC_a_*s or via de-differentation of TAC cells back into stem cells. As such the *SC_a_* loss must be viewed in the context of *irreparable injury*. From our analysis of the homeostatic scenario we know that there is a direct relationship between the number of TACs and the number of *SC_a_*s at steady state: ablating *X*% of the *SC_a_* population will decrease the steady state distribution of TAC cells by *X*%, and we investigate the *re-establishment time* required to reach the new distribution. Specifically, we calculate

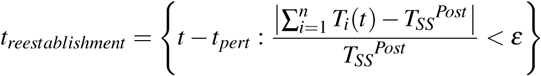

where *ε* is arbitrarily small, *T_SS_^Post^* is the total number of TAC cells that would be generated at the new steady state after *X*% of initial stem cell population is removed, and *t_pert_* is the time the stem cell loss perturbation occurs. Before moving to the perturbation experiments, it is sensible to first investigate the impact of *ε* for the re-establishment time. To do so, we perform Gillespie-based stochastic simulations for the specific parameter set described in Subsection 4.1, under 3 different values of *ε*: *ε* =0.01,0.05,0.1 given a 20% *SC_a_* loss. Value of *ε* =0.01,0.05,0.1 represent reaching within 1%, 5%, 10% of the new TAC population homeostatic state. Inevitably, when *ε* is very small, the system is in a noisy regime and it is unfeasible to get exactly within *X*% of the new TAC population level after applying the perturbation. In this case there can be large variations in the re-establishment time. Hence, we focus on finding a value of *ε* which is small enough to get close to the new level, but not small enough that we start to face sensitivity issues. Our results (see Figure 9a) show that if we impose a very tight bound restriction (*ε* = 0.01) then large variations in the re-establishment time are possible. On the other hand, for *ε* = 0.05 or 0.1, variations in *t_reestablishment_* are relatively small, indicating values *ε* ≥ 0.05 are required and we choose *ε* = 0.05. Moreover, for *ε* = 0.05 both the mean of the stochastic model and the deterministic model agree (see Figure 9a for *ε* = 0.05 and Figure 9b for 20% *SC_a_* loss) therefore allowing us to use the deterministic model to explore the mean re-establishment time for different perturbations.

**Fig. 9:**
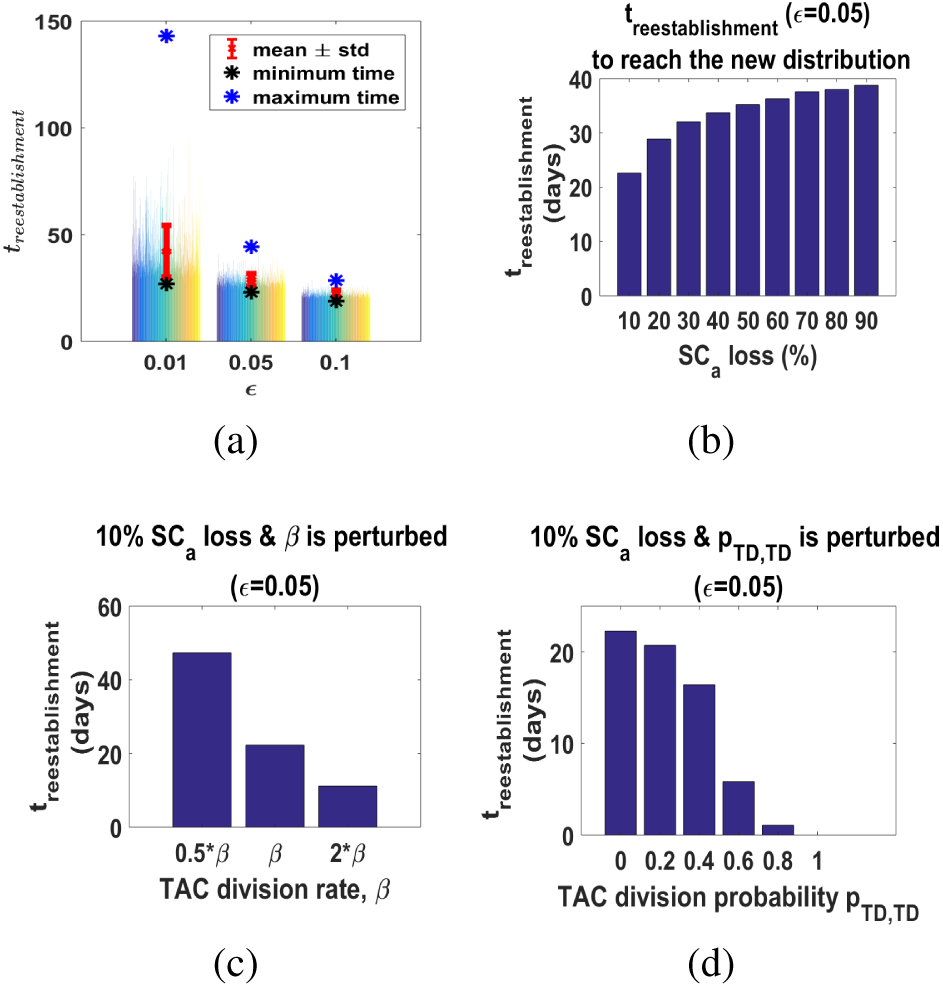
(a) Each line represents the computed re-establishment time for an individual stochastic simulation (total of 1,000 simulations) under 3 different values of *ε*. Also plotted are the mean, standard deviation, maximum and minimum re-establishment time (in days), using the reference parameter set. Note that each coloured line in each bar represents the result from one stochastic simulation of the model, however only a subset of the total simulations is plotted for each case. The specific simulation that generates one of the extrema (maximum or minimum) may therefore not be represented. (b)-(f) Plots show different scenarios for a mouse corneal epithelium’s fate resulting from pathological scenarios, with *ε* = 0.05. (b) Required time to reach the new steady state according to % loss in *SC_a_* number. (c) Case of 10% *SC_a_* loss. An increase in *β* decreases the time needed for the epithelium to collapse. (d) Case of 10% *SC_a_* loss. An increase in *p_TD,TD_* decreases the time needed for the epithelium to collapse.

In Figure 9b we show that large increases in *SC_a_* loss only result in moderate increases in the reestablishment time. Specifically, when 10% of the *SC_a_* population is lost, *t_reestablishment_* ≈ 23 days. Moreover, when 90% of *SC_a_*s are lost then the epithelium reaches its new steady state in roughly 39 days. Thus, despite the 9 fold difference in *SC_a_* loss, there is a less than 2 fold increase in the time required. Hence, from a clinical/biological perspective, in the case of a patient who suffers from sudden limbal SC loss, clinicians could be able to predict the time at which the new homeostatic state is attained.

To understand how parameter changes alter the reestablishment time, we perturb TAC, SCa division rates and probabilities. In Figure 9c we show the impact of ×2 and ×1/2 perturbations to the default value of *β* on the re-establishment time 9b: clearly, *β* has significant impact with ×2/ × 1/2 perturbations generating corresponding-sized perturbations on re-establishment time. On the other hand, equivalent simulations involving perturbations to *α* show no effect on *t_reestablishment_* (data not shown). These results can be anticipated by the analytical solution to the ODE system, Equation 13, where we see the intrinsic link between *β* and *t*; *α*, on the other hand, simply enters via a scaling. Assuming now that under a biological experiment, in addition to the 10% *SC_a_* loss, TACs are forced into premature TD differentiation. An increase in probability *p_TD,TD_* will result in epithelium collapse in shorter time (see Figure 9d). Note that no substantial change was observed for increases to *p_T,TD_* while keeping *p_TD,TD_* constant, due to the insubstantial difference in the number of TACs lost.

### 4.3 TAC Loss Perturbation Experiments

Here, we investigate the capacity of the epithelium to recover under insults to the central epithelium area: perturbations to the TAC population from their homeostatic (steady state) values. We assume perturbations do not change the number of *SC_a_*s, and therefore do not expect any change to the homeostatic situation post recovery. The *recovery time* is defined as the time it takes before returning to the pre-perturbation TAC number:
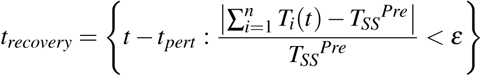

where *ε* is arbitrarily small, *T_SS_^Pre^* is the total number of TAC cells before the perturbation is applied. As in the previous subsection (see Subsection 4.2), we investigate the impact of *ε* for the recovery time. Similarly to the *SC_a_* perturbation experiments we found that *ε* = 0.05 represents a suitable small value without facing sensitivity issues. The deterministic model captures the mean recovery time of the stochastic model, that is *t_recovery_* = 39.7 days (see Figure 10a for *ε* = 0.05 and Figure 10b for remaining 0% of the TAC population) and, hence, in the rest of the section we exploit the deterministic model to explore *t_recovery_* under different perturbation experiments.

**Fig. 10:**
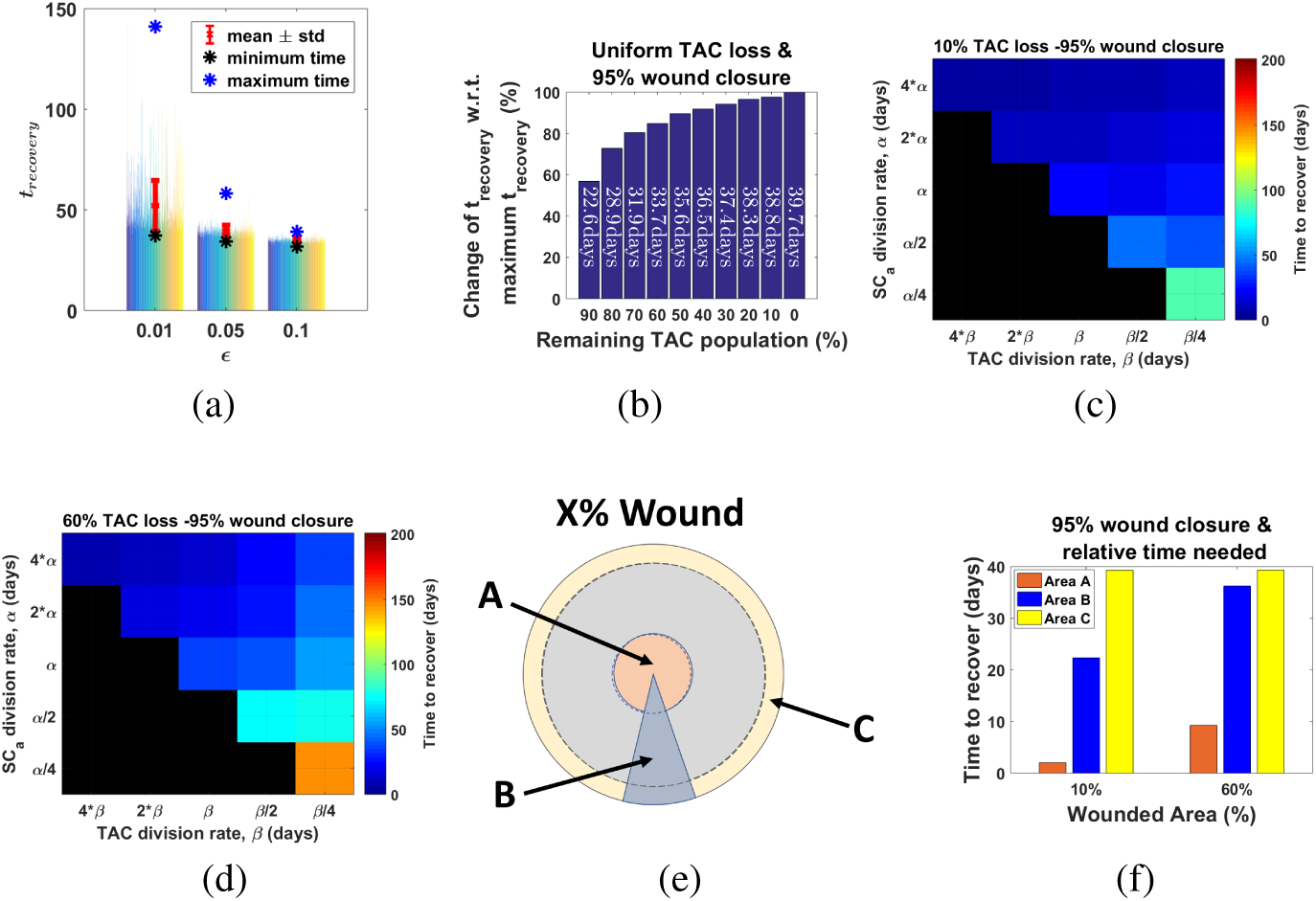
(a) Each line represents the computed recovery time for an individual stochastic simulation (total of 1,000 simulations) under 3 different values of *ε*. Also plotted are the mean, standard deviation, maximum and minimum recovery time (in days), using the reference parameter set. Note that each coloured line in each bar represents the result from one stochastic simulation of the model, however only a subset of the total simulations is plotted for each case. The specific simulation that generates one of the extrema (maximum or minimum) may therefore not be represented. (b)-(f) Plots show different scenarios for a mouse corneal epithelium’s fate following a wound type perturbation, with *ε* = 0.05. Specifically: (b) The percentage change of recovery time with respect to the maximum recovery time, assuming uniform removal across TAC generations. The required time for recovery is counted in days (in red). (c)-(d) Measurement of recovery time when TACs are lost uniformly across TAC generations and division rates *α* and *β* are perturbed. Black area correspond to no full recovery within 50 day period. (e) Perturbed areas: Area A, TAC loss weighted to higher generations; Area B, uniform TAC loss across all generations; Area C, weighted to lower generations. (f) Comparison between a minor (10%) and a large (60%) TAC loss according to the time needed for 95% recovery and the perturbed area.

We first consider *uniform perturbations*, where we remove equal percentages of each TAC generation. The impact on recovery time is summarised in Figure 10b which shows the *percentage change of recovery time with respect to the maximum recovery time* (i.e. the recovery time if 100% of TAC population was removed). As could be expected, increased TAC loss demands an increase in recovery time, although the results show that the change is relatively small with respect to the size of perturbation.

There is likely to be uncertainty in parameters such as *SC_a_* and TAC division rates, and particularly whether they change in the face of some perturbation (cell proliferation rates can be experimentally manipulated e.g. Lehrer et al. 1998, Saghizadeh et al. 2017. Hence, we vary these parameter values and plot the resulting change to the recovery time in Figures 10c-10d. Decreasing the *SC_a_* division rates (*α*) has a negative effect on recovery, and can even lead to recovery failure as alterations of this type act to lower the homeostasis level of cells. Similarly, decreases to TAC division rate will slow down the recovery rate (as expected from our earlier analytical solution, Equation 13). Thus, in the context of perturbations to TAC numbers, a (possibly temporary) response of decreased TAC and increased *SC_a_* division rate would be optimal for quick recovery, in line with certain biological findings (Lehrer et al., 1998; Pal-Ghosh et al., 2004). Note that, as expected, recovery time increases according to the size of TAC loss, cf. Figures 10c and Figure 10d.

We next investigate the effect of non-uniform perturbations, removing a set percentage of the total TAC population but the removal weighted variably across the TAC generations. TAC generations are assumed to be distributed radially, since cells move centripetally (Nagasaki and Zhao, 2003) over time, and thus different weightings in this manner would correspond to principally removing the TAC cells from the centre or the periphery of the cornea. Figure 10e illustrates our three basic perturbation types: a type A perturbation corresponds to predominantly removing higher generation TAC cells (expected to be located in the central cornea region); a type B perturbation corresponds to an equal weighting removal across all generations; a type C perturbation corresponds to predominantly removing lower generation TAC cells (expected to be located in peripheral regions). Note that for perturbation A (C) we first removed all cells from the highest (lowest) generation, followed by the next higher (lowest) generation and so forth until the required number has been removed. As Figure 10f suggests, area A perturbations show significantly faster recovery rates than rate C perturbations, suggesting that preserving first generation TAC cells is more critical for recovery: intuitively, low generation cells and their descendants can remain in the basal layer for significantly longer before automatic terminal differentiation. In fact, for an area C perturbation we even see a drop in the total number of TACs in the initial stages of the recovery process. Summarising, we expect that the region where a perturbation is applied may have some relatively significant impact on the subsequent recovery time.

## 5 Conclusion

In this paper, we have developed a purposefully simple stochastic mathematical model, based on an analogy to chemical reactions, to clarify the main factors involved in maintaining the corneal epithelium. We have focused on the proliferation process of both *SC_a_*s and TACs, considering only the dynamics in the basal epithelial layer and thereby assuming that maintaining this layer provides the key to epithelial homeostasis.

Our analysis provides an explicit link between the number of TACs at each generation and: (i) the numbers of active stem cells, and (ii) the relative rate of *SC_a_* to TAC division. Further, the TAC proliferation rate who has a significant impact on the rate of temporal dynamics of TACs. For the TAC division probabilities, we considered two potential forms: (i) an analytically convenient step-function form, and (ii) an exponential form. We have shown that these two reasonably plausible forms give very similar results and hence, there is robustness of the results with respect to the precise form. The analysis of the model using these probability functions generated parameter spaces for the constraints under which the epithelium maintains.

To account for the variability about the mean TAC proliferation process, and hence investigate the noisiness of the system, we derived the second moments at the steady state using the Lyapunov equation and then calculated the FF and CV for each TAC generation. The work on Fano Factor and Coefficient of Variation presented here suggests that an evolving less noisy system might be fitter to avoid a noisy behaviour of cells and hence maintain the epithelium. We further investigated the required number of TAC generations to maintain the corneal epithelium by letting *t_TAC_* range across the acceptable range of TAC proliferation rates and 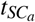 vary widely. For a mouse corneal epithelium we found that when *SC_a_*s divide once every 14 days, which is assumed to be towards the upper bound of the *SC_a_* proliferation rate, the required TAC generation number for the epithelium maintenance increases, although not much compared with a 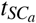 of 2 days. Nevertheless, the 14 days figure may be because SCs switch in and out of the active state. Thus, the 14 days could be viewed as more appropriate for the total stem cell population (rather than the active population). Further, we are able to make a direct comparison with respect to the number of the required TAC generations between species. For example, we can compare the mouse with the rabbit corneal epithelium. The rabbit corneal epithelial area is bigger and TACs proliferate faster than in mouse (once every 18 hours). This would imply that 5 more TAC generations will be needed for maintaining the rabbit corneal epithelium, compared to those needed for the mouse, assuming that the *SC_a_* division rate remains the same.

The work in this paper serves as a stepping stone in understanding the maintaining process of the corneal epithelium and its behaviour under perturbations linked to pathological/wound healing type scenarios or biological experiments. Of course, cell migration is fundamentally important but has been neglected here in order to concentrate on the proliferation kinetics of cells. Future work will use both PDE and random walk description of motile cells (Grima and Newman, 2004; Grima, 2008) to extend the present model into a spatial one capable of modeling the centripetal movement of TAC population seen in biological experiments, and obtain a clearer idea of corneal epithelium wound healing responses. Other possible extensions will be to include the quiescent SC population to allow some feedback mechanisms.

## Appendix A Parameter Estimations

### A.1 Mouse

We first note that the corneal circumference of a mouse is ~ 10,000*µm* (Di Girolamo et al., 2015; Dorà et al., 2015) and a typical basal cell diameter is ~ 10*µm* (Romano et al., 2003). If stem cells simply formed a one-cell thick ring, a total stem cell population of ~ 1,000 cells could be accommodated along the corneal-limbal border. Note, however, that an estimated 250 − 300 are active (Dorà et al., 2015) at homeostasis. To accommodate scenarios that can range from healthy to pathological, or eye sizes from larger to smaller, we assume the number of SCa in the limbus ranges between 100 − 1000.

Although the cornea is dome-shaped, for the purposes of the model we have assumed it is a hemisphere with a circumference of approximately 10,000 *µm*. Then, the radius of the corneal is *r_corneal_* = 1,592 *µm*, from which the corneal area is 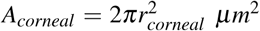. Similarly, the average area occupied by a basal corneal cell (assuming that the cell is a disc in the 2*D* plane) is 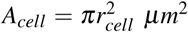, where *r_cell_* = 5 *µm*. Thus, an estimate of cells that can fit in the corneal epithelium is given by:
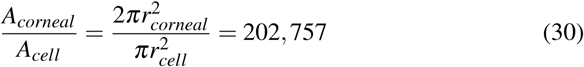

and to take into account not just the normal conditions, we can introduce the magnitude of 10^5^ as a guideline baseline value for the number of cells required to populate a small cornea.

For mouse we have a number of sources that provide indications of stem cell and TAC division rates. If it is assumed that mouse limbal epithelial SCs are equivalent to BrdU “label-retaining cells”, which include slow-cycling stem cells, it can be estimated that *certain* limbal epithelial SCs do not divide more often than once per two weeks (~ 14 days). This calculation follows from detectable BrdU retention for at least 10 weeks (Douvaras et al., 2013), and that BrdU is probably diluted to undetectable levels after 4 - 5 cell divisions (Wilson et al., 2008). However, this is quite likely to provide an approximate lower bound for division rates, as it remains quite possible that certain SCs divide significantly more quickly and may not be detected by the label-retaining cell approach. As such, the mean SC cell cycle time may be considerably less than 2 weeks. Of course division rates are ultimately bounded by the minimum length of time needed to complete the cell cycle, which would be of the order of several hours to a day. Consequently, we take a range 6 hours to 16 days for (active) stem cell doubling times.

Experimental studies on the TAC cell cycle in the peripheral corneal epithelium indicate that almost 50% of basal corneal epithelial cells are in S-phase of the cell cycle, during a 24-hour labelling period (Urbanowicz et al., 2011). This suggests a minimum cell doubling time of just over 2 days but it would be longer if certain TACs cycle more slowly. Similarly, an average mitotic rate of 37% of basal layer cells per day can be derived for rats from the results reported by Bertalanffy and Lau (1962) and this suggests a minimum cell doubling time of about 2.7 days. (The original results showed that 14.5% of all corneal epithelial cells divided per day and results for the mouse imply that about 38.8% of mouse corneal epithelial cells are in the basal layer (Douvaras et al., 2013)). Other experiments on the TAC cell cycle in the peripheral corneal epithelium have estimated it as approximately as 72 hours for the mouse (Lehrer et al., 1998). Overall the results show that the average doubling time for TACs is about once every 2 - 3 days but may be longer in the central corneal epithelium (Lehrer et al., 1998). While we centre on an average rate of 2 days, for our studies we again use a range of 6 hours to 16 days to include scenarios under normal and abnormal conditions.

### A.2 Human

Experimental data suggests that the average corneal diameter in human eye is 11.71 ± 0.42 mm, (Rüfer et al., 2005) implying a corneal circumference ~ 36.770 *mm*^2^. In the absence of specific data, we consider an analogous case to the mouse and suppose the circumference corresponds to the corneal-limbal border. Assuming limbal corneal cells are 10*µm* in diameter, we estimate that there is a room for ~ 3,000 − 4,0000 limbal cells forming a one-cell thick ring; although (in contrast to the mouse case) some biological studies suggest that they are asymmetrically distributed (Wiley et al., 1991; Pellegrini et al., 1999; Shanmuganathan et al., 2007). If a similar fraction (to that of mouse) of this population is taken to be active, we estimate ~ 1,000 active stem cells (*SC_a_*) in the human limbus. Again, we consider an order of magnitude range about this value (~ 400 - 4,000).

Using the same calculations adapted from the mouse case gives an order of magnitude of 10^6^ basal epithelial cells fitting in the human cornea.

### A.3 Rat and Rabbit

To demonstrate variability across other species, we note that rat and rabbit corneas have average diameters of 5.5*µm* (Cabrera et al., 1999) and 14.375*µm* (Tsonis, 2011) respectively. Straightforward calculations show that the circumferences will be 17, 270*µm* and 45, 138*µm* respectively. Making the same assumptions as earlier, this would allow for a total of 1, 727 and 4, 513 stem cells and, if again approximately 1/4 are active, ~ 450 and ~ 1200 active stem cells for rat and rabbit respectively. Calculating an estimate for the total number of cells that can fit into the basal epithelium yields a magnitude ~ 10^5^ for rat and ~ 10^6^ for rabbit, the former the same magnitude as the mouse and the latter similar to the human eye.

For a rabbit corneal epithelium, experimental data on TAC doubling time suggests once every 18 hours (3/4) (Castro-Muñozledo, 1994). We are lacking such data for the rat eye. Nevertheless, the parameter spaces provided throughout the paper can give a rough estimate of the TAC generations required for the epithelium maintenance for each of rat and rabbit eye.

## Appendix B Derivation of Matrices Included in Lyapunov Equation

### B.1 Jacobian Matrix

The Jacobian matrix **J** can be derived from the stochastic mean system (5)-(7) obtained in Section 2.3. Matrix **J** of our *n*-ODEs system for the stochastic means of

TACs is:
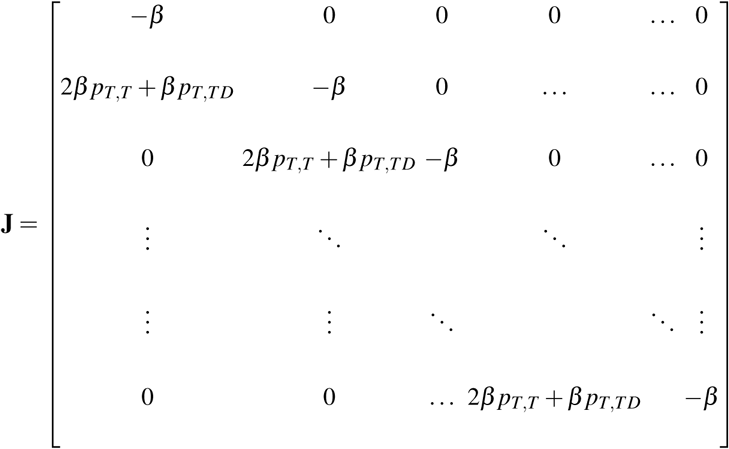

Note that 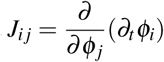 where *φ_i_* = *N_Ti_* and *φ_j_* = *N_Tj_* with *j* = 1, …, *n*.

### B.2 Stoichiometric Matrix

For the stoichiometric matrix we are only interested in the number of TACs at each reaction. Denoting the reactions as *r_k,l_* with *k* the reacting population (i.e. *k* = 0, 1, ·*, n*, where *k* = 0 corresponds to the *SC_a_* division to *TAC*_1_ and *k* = *i* the *TAC_i_* divisions) and the pathway indicator is *l* (hence *l* = 1, 2, 3). The reactions can be written as
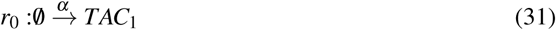

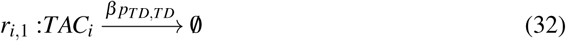

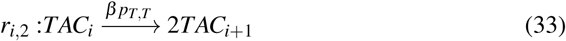

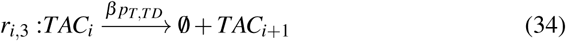

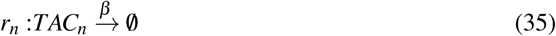

where 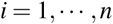 to be the number of TAC generation. The stoichiometric vector for *TAC*_1_ is 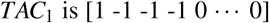, for *TAC_i_* is 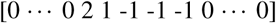 and for *TAC_n_* is 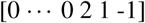. As an example for the stoichiometric matrix **S**, let us assume that the total number of TAC generations is 3, then
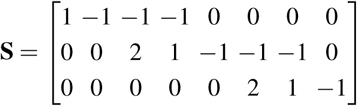

Note that in the stoichiometric matrix for *n* TAC generations, the number of zero elements at the start of each row (excluding the first and last row which correspond to the first and last TAC generation respectively) will be *N*_0_ = 3(*i* − 1) + 2 with *i* = 2, …, *n* − 1. Hence, the position of the first non-zero element in each row (*i* = 2, … *, n* − 1) follows the sequence 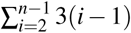.

### B.3 Vector of Macroscopic Rates

To find the vector of macroscopic rates 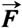 we recall the reactions 31–35 listed in Appendix B.2 with corresponding rates:
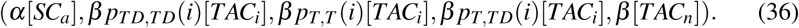

Hence, the vector 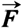 for our system is
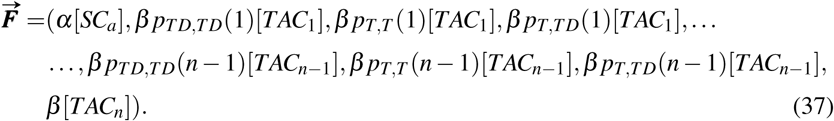

### B.4 Diffusion Matrix

For the elements of the diffusion matrix, as already discussed in the text, we used
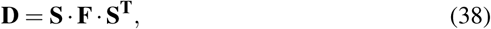

inside the matlab code, with **F**, **S** and **S^T^** determined as above.

## Acknowledgements

The authors would like to thank Dr. John D. West (University of Edinburgh) for his valuable help in understanding the underlying mechanisms of the corneal epithelial maintaining process and the data provided. Eleni Moraki was supported by The Maxwell Institute Graduate School in Analysis and its Applications, a Centre for Doctoral Training funded by the UK Engineering and Physical Sciences Research Council (grant EP/L016508/01), the Scottish Funding Council, Heriot-Watt University and the University of Edinburgh. Ramon Grima would like to acknowledge funding from BBSRC grant BB/M025551/1. Kevin J. Painter would like to acknowledge Politecnico di Torino for a Visiting Professor position and funding from BBSRC grant BB/J015940/1.

